## Supplemental material legends for "Decoupling of mRNA and protein expression in aging brains reveals the age-dependent adaptation of specific gene subsets"

### Supplemental Figure Legends

**Supplemental Figure 1. Anatomical characteristics of the aging brain.**

**(A)** Representative coronal cuts of 6-month (6M) and 24-month (24M) whole brains stained with hematoxylin and eosin. Scale bar: 1mm. **(B)** The total cerebral cortex area of 6-month-old (grey) and 24-month-old (red) mouse brains (n=8 per age group; 4 males and 4 females) was measured. Results are indicated as mean ± SD. ns: not significant **(C)** Graph representing the weight of each mouse before sacrifice. The 6-month-old mice are in grey (n=61; 42 males and 19 females), and the 24-month-old mice in red (n=59; 40 males and 19 females). (**) $p < 0.0001$ in one-way ANOVA; ns: not significant.

**Supplemental Figure 2. Location of differentially expressed genes on chromosomes.**

The genes whose expression is upregulated **(A)** and downregulated **(B)** in the cortex during aging are represented by red dots. The purple lines indicate regions where these genes are statistically enriched, compared to the density of genes in the background. The genome was scanned with a sliding window of 6 Mb. Within each window, a hypergeometric test was used to determine if your genes are significantly overrepresented (window FDR cutoff: 0,001).

**Supplemental Figure 3. Correlation between transcript and protein abundance.**

**(A)** Concentration of proteins extracted from 6-month-old (6M) and 24-month-old (24M) cortices. (n=6 per age group; 3 males and 3 females). Results are indicated as mean ± SD. **(B)** Venn diagram showing the overlap between differentially expressed genes (DEG) (n=427) and proteins identified by mass spectrometry (n=2,413) in the aging cortex. On the right, the list of 13 DEG (3 upregulated in red, 10 downregulated in blue) detected by mass spectrometry. **(C)** Histogram of the number of detected peptides per gene, ranked based on the gene TPM value. The 10,000 top genes are presented. TPM: transcripts per million. **(D)** Heatmap of the gene expression (in TPM; transcript per million) for genes for which the protein was found differentially expressed during aging (in black; n=206) and genes for which the transcript was found differentially expressed during aging, but the protein was not detected (in gray; n=427). The counts were normalized per column, the most expressed gene in one sample having the same score as the most expressed in another sample (n=3 per age group). **(E)** Workflow of the proteomic analysis.

**Supplemental Figure 4. Controls and originals from Figure 4.**

**(A)** Protein levels of GAP43 within the 6- and 24-month-old cortex were analyzed by western-blot. Actin was used as a loading control. Molecular weight markers of the ladder are indicated on the side. On the left panel, original images used for the Figure 5A. Red frames indicate the image sections selected for the figure. On the right panel, original images used to quantify GAP43 signal normalized by actin signal (n=4 per age group). **(B)** GAP43 immunofluorescence on coronal cuts of 6-month-old brain. Images illustrate the representative staining in the cortex motor area with the GAP-43 antibody (green) and DAPI (blue). At the top panel, immunofluorescence without a GAP43 primary antibody and at the bottom panel, immunofluorescence with a GAP43 antibody. Scale bar: 100 μm.

### Supplemental Table Legends

**Supplemental Table 1. Table of RNA-seq count data.** The table displays the expression of 55,421 genes in the 6- and 24-month-old cortices (respectively Cx-6 and Cx-24) (n=3 per age group). Values for each gene are in TPM (transcripts per million). The “Heatmap” column indicates whether the gene is present in the heatmap of Fig. 2C.

**Supplemental Table 2. List of genes differential expressed in the aging cortex.** The table displays 427 genes differentially expressed in the 6- and 24-month-old cortices (respectively Cx-6 and Cx-24) (n=3 per age group). Values are in log(CPM) (counts per million) and normalized for each gene. The “rank” column representing the level of mRNA expression was calculated from the RNA-seq count data (Supplemental table 1), and ranges from 1 to 55,421.

**Supplemental Table 3. List of proteins detected by mass spectrometry in the aging cortex.** The table (MaxQuant output) displays 2,471 proteins detected by mass spectrometry in the 6- and 24-month-old cortices (respectively Cx-6 and Cx-24) (n=2 per age group). Several sheets with the different basic filtration steps are included in the table.

**Supplemental Table 4. Data analysis table of proteins identified in the aging cortex.** The table displays proteins detected in the 6- and 24-month-old cortices (respectively Cx6 and Cx24) (n=2 per age group). The “quantitative data” sheet compiles the LFQ (label-free quantitation) intensity values for the 740 proteins obtained after filtration. The “feature meta data” sheet displays all the mass spectrometry information including the peptide number, the coverage and the LFQ intensity.

**Supplemental Table 5. List of proteins differential enriched in the aging cortex.** DOWN: proteins downregulated in the cortex during aging (n=42). UP: proteins upregulated in the cortex during aging (n=164).

**Supplemental Table 6. Selected peptides for PRM mass spectrometry analysis.**
