## Supplementary figures and images for "Decoupling of mRNA and protein expression in aging brains reveals the age-dependent adaptation of specific gene subsets"

### Supplemental Figures

#### Supplemental Figure 1

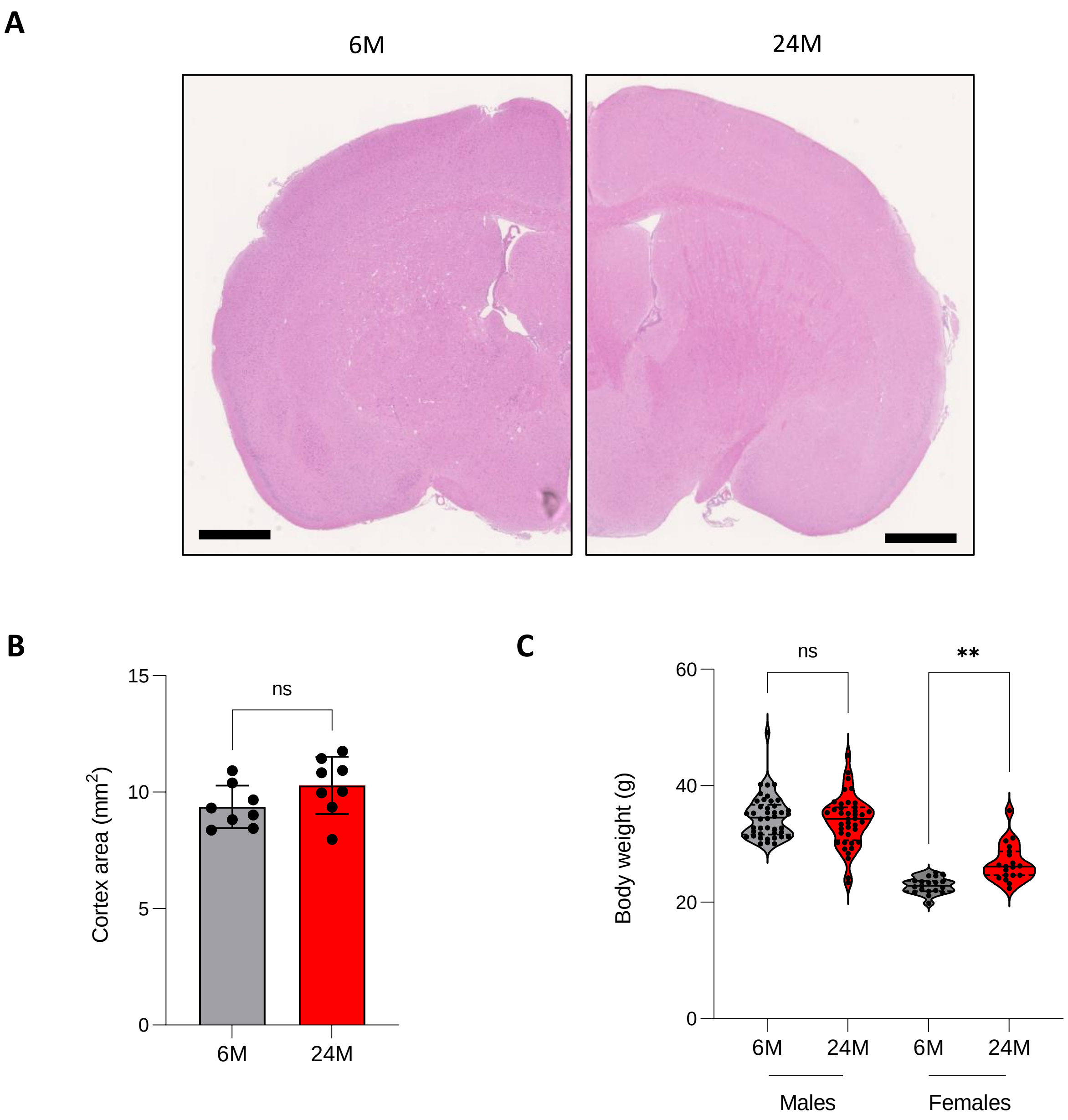

#### Supplemental Figure 2

**A**

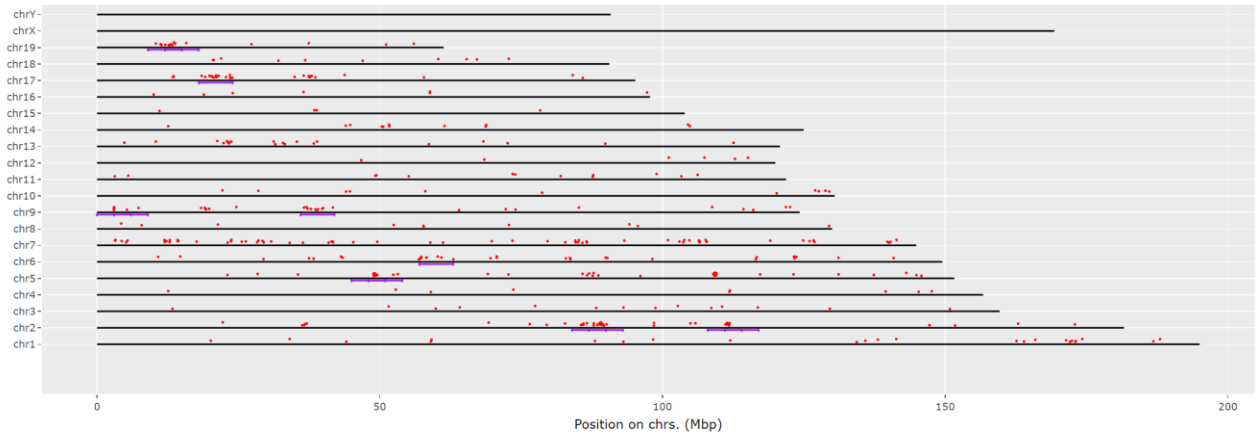


**B**

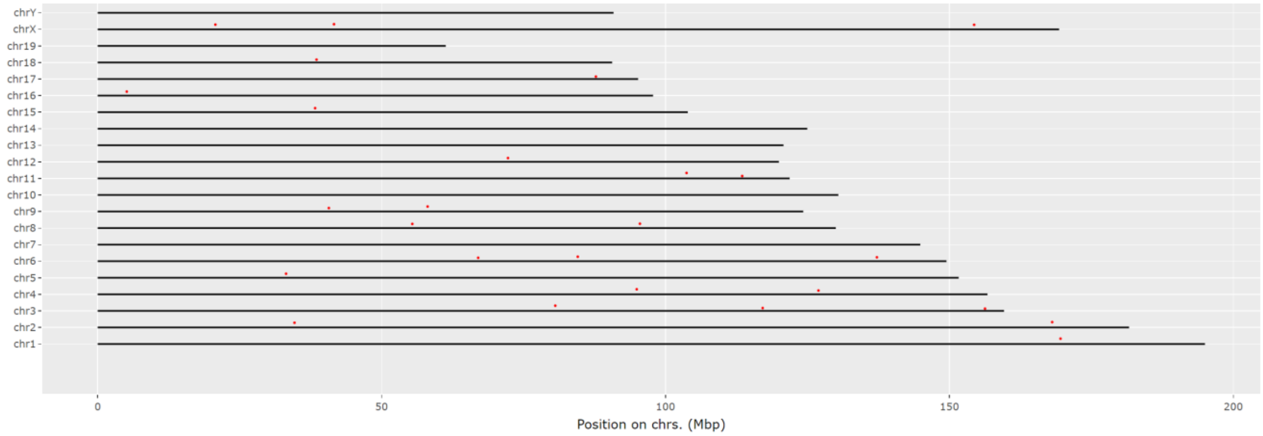

#### Supplemental Figure 3

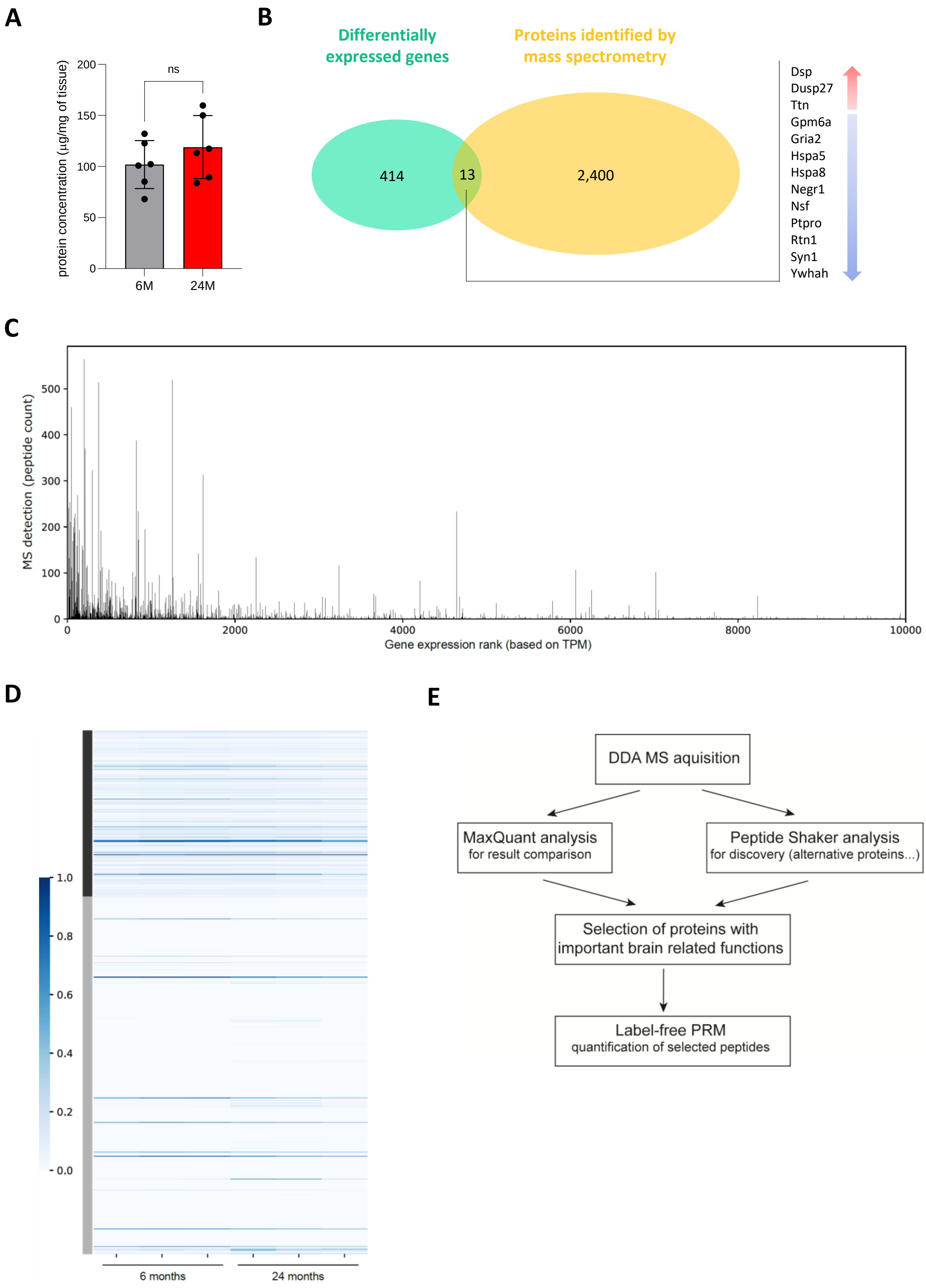

#### Supplemental Figure 4

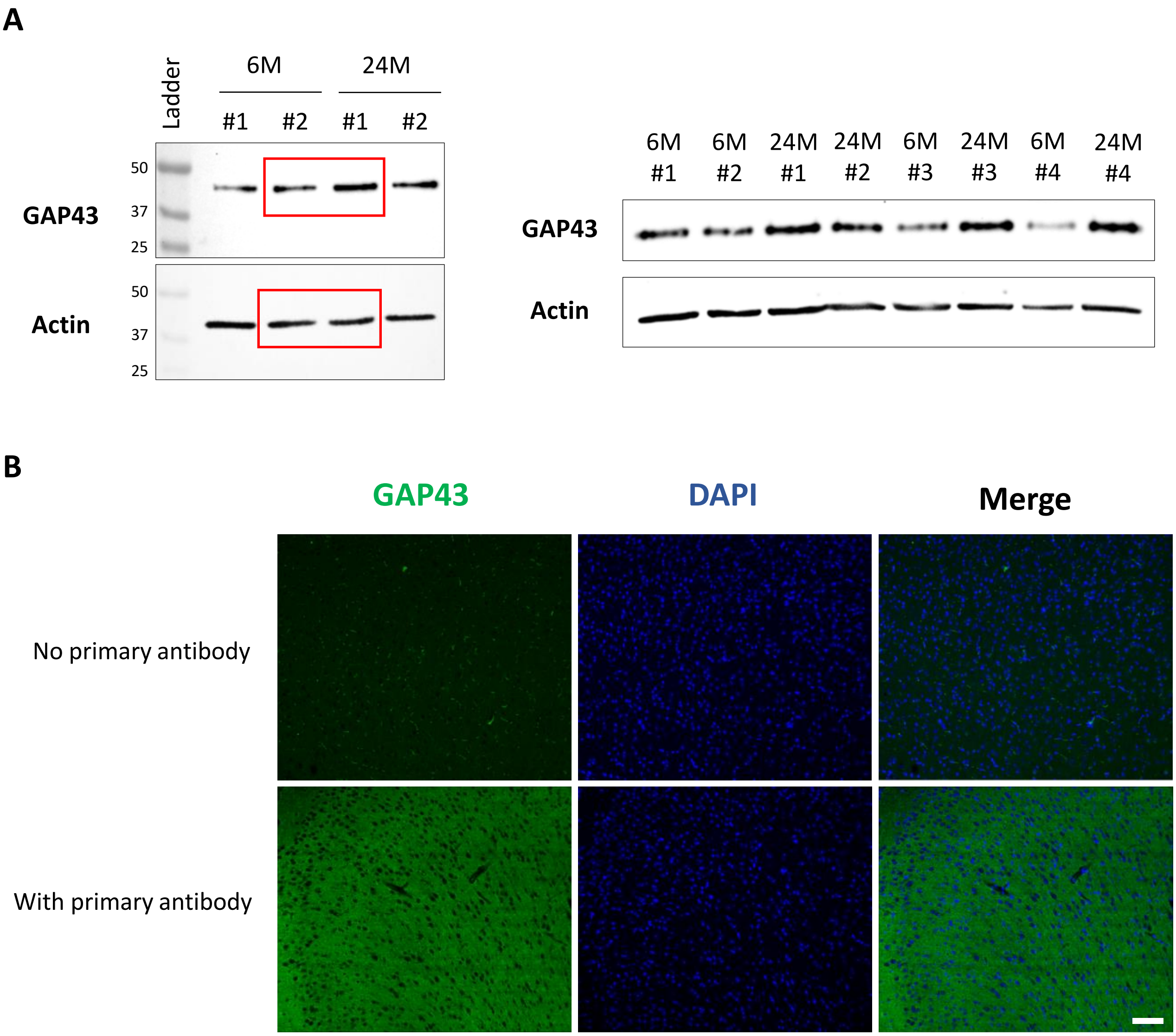
