## Supplemental figure 5 for "Decoupling of mRNA and protein expression in aging brains reveals the age-dependent adaptation of specific gene subsets"

### Supplemental Table 4

| DOWN | | UP | | | | | | | |
|---|---|---|---|---|---|---|---|---|---|
| Apoa1 | Pdcd5 | Abi1 | Atcay | Dlgap1 | Gad1 | Lancl2 | Pafah1b1 | Psmc1 | Strap |
| Apoo | Pin1 | Abi2 | Atl1 | Dmtn | Gad2 | Lmnb2 | Pak1 | Psmc4 | Tardbp |
| Arf5 | Ppa2 | Acaa2 | Bcat1 | Dnaja2 | Gap43 | Lrrc47 | Pcbp2 | Psmc5 | Tceal5 |
| Arl8a | Rpl18 | Acad9 | Bcs1l | Dnajc11 | Gps1 | Lsamp | Pcca | Psmd11 | Tmx3 |
| C1ql3 | Rpl19 | Acadl | Bpnt1 | Dpysl5 | Gsk3a | M6pr | Pccb | Psmd13 | Tmx4 |
| Cnksr2 | Rpl23a | Acadm | Cacnb3 | Dync1li1 | Gucy1b3 | Maob | Pcyox1l | Psmd3 | Tom1l2 |
| Cplx2 | Rpl38 | Acadvl | Cadm4 | Dync1li2 | H2afy | Mapk3 | Pde1b | Psmd5 | Tomm40 |
| Csrp1 | Rpl7a | Acot9 | Camk2d | Eif3e | Hapln1 | Mapre2 | Pdia6 | Psmd6 | Tuba8 |
| Dctn3 | Rpl8 | Acsl1 | Cct5 | Eif4a1 | Hars | Me3 | Pepd | Ptpn9 | Txnrd1 |
| Diras1 | Rpl9 | Actr1b | Cct6a | Eif4a3 | Hebp1 | Mecp2 | Pick1 | Rars | Uba3 |
| Dynll2 | Rps10 | Adk | Cct7 | Elavl4 | Hexb | Mtch1 | Pip4k2a | Rmdn3 | Upf1 |
| Enoph1 | Rps26 | Adrbk1 | Cd200 | Entpd2 | Hnrnpl | Nadk2 | Pip4k2c | Rraga | Vat1l |
| Fabp5 | Stmn1 | Ahcy | Cd47 | Epm2aip1 | Hnrnpm | Nap1l4 | Pls3 | RtcA | Vim |
| Fam162a | Stmn2 | Ak5 | Cmpk2 | Erlin2 | Homer2 | Nisch | Pmpca | Ruvbl1 | Vps26b |
| Gng2 | Timm9 | Aldh6a1 | Cndp2 | Etf1 | Hspa2 | Nova1 | Ppm1f | Scrn1 | Vps45 |
| H2afv | Tmem11 | Aldh7a1 | Coro2b | Etfdh | Kbtbd11 | Nsfl1c | Ppp2r5a | Sh3glb1 | Wasf3 |
| Lin7b | Tpm4 | Anxa3 | Csde1 | Fam126b | Kctd12 | Nt5dc3 | Ppp2r5c | Shroom2 | Yars |
| Lin7c | Ttr | Ap3m2 | Cyp46a1 | Farsa | Kctd16 | Ola1 | Ppp5c | Slc9a3r1 | |
| Ndufa7 | Txn | Arfip2 | Dars | Fdps | Kiaa0513 | P4hb | Prkar1a | Smap1 | |
| Nme2 | Uchl3 | Ass1 | Dbnl | Flot2 | Klc2 | Pa2g4 | Prkar2a | Snx5 | |
| Pdap1 | Uqcr10 | Atad3 | Dctn4 | Gabra1 | Lactb | Pabpc1 | Prmt8 | Src | |
