## Supplemental figure 6 for "Decoupling of mRNA and protein expression in aging brains reveals the age-dependent adaptation of specific gene subsets"

### Supplemental Table 5

| Gene | Peptide sequence | m/z | charge state | retention time |
|---|---|---|---|---|
| **Aldh2** | GYFIQPTVFGDVK | 735,88 | 2+ | 8464,1 |
| **Aldh2** | YGLAAAVFTK | 520,79 | 2+ | 6045,2 |
| **Apoa1** | VAPLGAELQESAR | 670,86 | 2+ | 4884,8 |
| **Apoa1** | SNPTLNEYHTR | 444,55 | 3+ | 1328,1 |
| **Apoe** | LGPLVEQGR | 485,27 | 2+ | 2303,2 |
| **Apoe** | TANLGAGAAQPLRDR | 504,27 | 3+ | 3095,3 |
| **App** | THTHIVIPYR | 412,90 | 3+ | 2845,6 |
| **App** | VESLEQEAANER | 687,83 | 2+ | 3678,6 |
| **Atp2b2** | GIIDSTHTEQR | 419,55 | 3+ | 2145,4 |
| **Atp2b2** | NVFDGIFR | 484,25 | 2+ | 7882,2 |
| **Atp5b** | IGLFGGAGVGK | 488,29 | 2+ | 6290,2 |
| **Atp5b** | VVDLLAPYAK | 544,82 | 2+ | 7307,6 |
| **Atp5o** | LGNTQGIISAFSTIMSVHR | 683,36 | 3+ | 8294,1 |
| **Atp5o** | LVRPPVQVYGIEGR | 528,31 | 3+ | 5435,7 |
| **Cadm2** | DGAELPDPDR | 542,75 | 2+ | 3507,4 |
| **Cadm2** | GKPLPEPVLWTK | 455,60 | 3+ | 5175,3 |
| **Cap2** | SALFAQLNQGEAITK | 795,93 | 2+ | 6710,7 |
| **Cap2** | VEYQEDRNDLVISETELK | 727,36 | 3+ | 5763,9 |
| **Cntn1** | DGEYVVEVR | 533,27 | 2+ | 3556,5 |
| **Cntn1** | GPPGPPGGLR | 452,75 | 2+ | 2980,0 |
| **Glul** | ATSASSHLNK | 529,27 | 2+ | 1830,1 |
| **Glul** | LTGFHETSNINDFSAGVANR | 717,35 | 3+ | 5769,9 |
| **Got2** | ASAELALGENNEVLK | 779,41 | 2+ | 5847,2 |
| **Got2** | VGASFLQR | 439,25 | 2+ | 4295,1 |
| **Ndufb10** | PDSWDKDVYPEPPSR | 596,61 | 3+ | 3499,7 |
| **Ndufb10** | TPAPSPQTSLPNPITYLTK | 1013,55 | 2+ | 9237,9 |
| **Ndufs2** | AVTNMTLNFGPQHPAAHGVLR | 562,54 | 4+ | 4192,5 |
| **Ndufs2** | LLNIQPPPR | 524,32 | 2+ | 5219,7 |
| **phtf2** | QVLVTVMK | 459,28 | 2+ | 5875,9 |
| **Prkcb** | IYIQAHIDR | 376,88 | 3+ | 2989,5 |
| **Prkcb** | LTDFNFLMVLGK | 699,38 | 2+ | 11666,8 |
| **Prrt2** | NSLQQGDVDGAQR | 694,33 | 2+ | 1519,3 |
| **Prrt2** | QEPASKPDVNR | 612,30 | 2+ | 1324,4 |
| **Psmd11** | TGQAAELGGLLK | 579,33 | 2+ | 5633,3 |
| **Psmd11** | YVRPFLNSISK | 441,92 | 3+ | 4118,2 |
| **Rap1gds1** | DLASAQLVQILHR | 488,62 | 3+ | 6796,0 |
| **Rap1gds1** | DQEVLLQTGR | 579,81 | 2+ | 3447,4 |
| **Scai** | HILELASILDVR | 460,27 | 3+ | 8878,7 |
| **Tomm70a** | AAAFEQLQK | 503,27 | 2+ | 3516,8 |
| **Tomm70a** | FALAQAQK | 438,75 | 2+ | 1955,3 |
| **Vim** | KVESLQEEIAFLK | 511,96 | 3+ | 6998,8 |
| **Vim** | QDVDNASLAR | 536,26 | 2+ | 4081,6 |
